# Further characterization and engineering of a 11-amino acid motif for enhancing recombinant protein expression

**DOI:** 10.1101/2024.10.14.618345

**Authors:** Jiawu Bi, Elaine Tiong, Weibiao Zhou, Fong Tian Wong

## Abstract

**Background:** Recombinant protein production in *Escherichia coli* (*E. coli*) is a widely used system in both academic and industrial research owing to its low cost and wide availability of genetic tools. Despite its advantages, this system still struggles with soluble expression of recombinant proteins. To address this, various solubility-enhancing and yield-improving methods such as the addition of fusion tags have been developed. However, traditional tags such as small ubiquitin-related modifier (SUMO) and Glutathione S-transferase (GST) can interfere with protein folding or require removal post-translation, which adds complexity and cost to production. To circumvent these issues, smaller solubility tags (<10 kDa) are preferred. This study specifically focuses on an 11-amino acid solubility-enhancing tag (NT11) derived from the N-terminal domain of a duplicated carbonic anhydrase from *Dunaliella* species.

**Results:** A comprehensive analysis was performed to improve the characteristics of the 11-amino acid tag. By investigating the alanine-scan library of NT11, we increased its activity and identified key residues for further development. Screening with the alanine mutant library consistently led to at least a two-fold improvement in protein yield for three different proteins. We also discovered that the NT11 tag is not limited to the N-terminal position and can function at either the N- or C-terminal of the protein, providing flexibility in designing protein expression constructs. With these new insights, we have successfully doubled the recombinant protein yields of valuable growth factors, such as fibroblast growth factor 2 (FGF2) and an originally low-yielding human epidermal growth factor (hEGF), in *E. coli*.

**Conclusion:** The further characterisation and development of the NT11 tag have provided valuable insights into the optimization process for such small tags and expanded our understanding of its potential applications. The ability of NT11 tag to be positioned at different locations within the protein construct without compromising its effectiveness to enhance recombinant protein yields, makes it a valuable tool across a diverse range of proteins. Collectively, these findings have the potential to simplify and enhance the efficiency of recombinant protein production.

## Background

*Escherichia coli* is commonly used for producing recombinant proteins in both academic and industrial settings. It is well-researched, grows quickly, and can be easily cultured in large quantities [1]. Protocols for using *E. coli* are cost-effective and offer a wide range of strains, reagents, promoters, and tools for producing functional proteins. While this recombinant protein system generally works well for microbial proteins, the protein translational environment within the microbial cells is often not suitable for various mammalian derived proteins, ranging from enzymes to growth factors. Research has shown that only 25% of human proteins can be expressed in *E. coli* in an active soluble form [2,3], with most proteins encountering difficulties during expression – resulting in insoluble aggregates or non-functional peptides [3,4]. While the simplicity of the bacterial system makes it easy to use, it presents challenges for expressing difficult proteins. Recombinant protein expression may fail due to improper disulfide bond formation, lack of necessary chaperones, or the host’s inability to perform required post-translational modifications [5,6].

To improve protein yields, gene fusion and solubility tags technology are often used. Traditional fusion solubility tags such as N utilization substance protein A (NusA), Glutathione S-transferase (GST), Maltose binding protein (MBP), and small ubiquitin-related modifier (SUMO) are employed to facilitate the folding and solubilization of recombinant proteins [7-11]. While these tags can improve the solubility and yield of fusion proteins [8-12], their large sizes may disrupt the folding and structural integrity of the fusion partners. This results in the production of soluble but non-functional or less active proteins [5,6,13], illustrating the limitations of these traditional solubility tags [7,12]. Therefore, this highlights the need to discover and develop new tags to complement existing ones, especially with the increasing use of microbial systems for protein expression.

More recently, shorter and smaller motifs have been discovered for enhancing solubility. One example is the Fh8 fusion tag (8 kDa) and its derivative, the H tag (first 11 amino acids of Fh8), which can not only serve as a purification tag, but also show high solubility, thermostability and increased protein expression in *E. coli* when attached to a protein partner. Compared to larger tags, smaller tags could potentially have larger scopes of application due to its inherent lower metabolic stress imposed on the host and reduced likelihood of interfering with the activity of the expressed protein [11,14]. Protein tags are now developed with more functionalities and flexibility in mind, offering novel solutions and expanding the design space for producing heterologous proteins in microbial systems.

In this study, we focus on a 11-amino acid protein motif, named NT11 [15], which can enhance protein solubility and yield in *E. coli* expression systems. NT11-tag was originally derived from the first 11 amino acid residues within the N-terminal N-half domain of a duplicated carbonic anhydrase (dCA) from *Dunaliella* algae species. The carbonic anhydrase enzyme was found to be composed of two predominant domains, a highly soluble N-terminal domain devoid of any enzymatic activity and a C-terminal domain with activity but on its own displayed lower expression levels and had poor solubility. Truncation experiments revealed the minimal peptide length of 11 residues, which was then adopted as a short acidic peptide tag capable of improving protein solubility and expression levels without interfering with protein structure or activity [15]. Previously we employed the tag to successfully solubilize and recover active proteins from a previously insoluble halogenase, as well as *in silico* mined PET-degrading enzymes [16,17].

Here, we conducted a detailed characterisation of the 11-amino acid motif to gain better insights and enhancement of the small but powerful tag. We first focused on examining the alanine-scan library of NT11 to enhance its activity and determine key residues for further engineering. Although the final optimized tag design varies depending on the protein of interest, screening with the alanine mutant library consistently resulted in at least a two-fold improvement in protein yield for three different proteins. Further investigation of NT11 also revealed new characteristics, where we demonstrate for the first time that its impact on enhancing protein production is not limited to a traditional N-terminal position; NT11 can function at either the N- or C-terminal of the protein of interest. This characteristic offers potential flexibility in designing the final protein expression constructs, allowing us to consider both solubility and purification aspects which ultimately expands tag functionality. Overall, further characterization and engineering of the NT11 tag have provided insights into the optimisation process of such small tags and expanded on the understanding of how the tag can be used. With these new insights, we have managed to enhance protein yields of high value growth factors, including fibroblast growth factor 2 (FGF2) and human epidermal growth factor (hEGF), in *E. coli*.

## Methods

### Construction of protein expression vectors and strains

Genes were codon-optimised for *E. coli* and synthesized from Twist Bioscience for assembly into pET28a (+) plasmids using *E. coli* OmniMax (Thermo Fisher Scientific, USA). *E. coli* XJb (DE3) (Zymo Research, USA) was used for protein expression. Amino acid sequences for the protein constructs are documented in Table S1.

### Mutagenesis

Mutagenesis of NT11 tag was created using the QuikChange™ Site-Directed Mutagenesis System developed by Strategene (La Jolla, CA), with short complementary primers (IDT, Singapore). The primers used for QuikChange™ are documented in Table S2.

### Protein expression

An initial rapid high-throughput screen of the NT11 alanine scan library was conducted with eGFP, LCC and Fast PETase. Single colonies (*E. coli* XJb (DE3)) were cultured in 1 mL of overnight autoinduction media (Merck, USA) supplemented with 50 µg/mL kanamycin, 1.5 M L-arabinose, and 0.5 M magnesium chloride. After 20 h of culturing at 28°C, the samples were pelleted at 21,000 g.

Experiments with LCC and brazzein used a 2-step process: 1) day 1 single colonies were inoculated into 1 mL Luria Bertani (LB) broth, containing 50ug/mL kanamycin (LB+ kanamycin), overnight at 37°C, then 2) equal densities were seeded into 50 mL LB+kanamycin, grown to 0.4-0.6 OD600, induced with 100 μM isopropyl-β-D-1- thiogalactopyranoside (IPTG), and cultured at 30°C for 20 h before pelleting at 4,000 g.

Growth factor expression used a day 1 LB with kanamycin starter culture, followed by day 2 inoculation into 1 mL overnight autoinduction media (Merck, USA) with 50 µg/mL kanamycin, 1.5 M L-arabinose, and 0.5 M magnesium chloride, cultured in a BioLector XT Microbioreactor (Beckman Coulter, USA) at 30°C, 1400 rpm, 25mL/min airflow for 20 h before pelleting at 21,000 g.

### Protein Purification

For all 1 mL protein expressions, the harvested cell pellets underwent a single freeze-thaw cycle in lysis buffer (50 mM sodium phosphate, 300 mM sodium chloride, 10 mM imidazole, 0.03% Triton X-100). The His-tagged proteins were purified via immobilized metal affinity chromatography (IMAC) using HisPur™ Ni-NTA resin (Thermo Scientific, USA), then eluted in 50 mM sodium phosphate, 300 mM sodium chloride, 500 mM imidazole.

For 50 mL protein expressions, the harvested pellets were resuspended in 10 mL lysis buffer, sonicated at 20% amplitude (10s/30s on/off, 5min40s), and clarified by 13,000 g centrifugation. The soluble His-tagged proteins were then captured on Ni-NTA resin and eluted in the same buffer as the 1 mL scale.

### Quantitation of Protein Expression

Purified proteins were eluted in volumes suited to their scale: 100 μL for 1 mL high-throughput, 2 mL for 50 mL shake flask, 1 mL for 1 mL BioLector. 10 μL of each elute was mixed with Laemmli Sample Buffer (Bio-Rad, USA), boiled at 95°C for 5 min, and run on SDS-PAGE gels (Thermo Fisher Scientific, USA). Gels were stained for ≥30 min with InstantBlue® Coomassie Protein Stain (Abcam, UK). Protein yields were quantified by densitometric analysis using ImageJ, normalizing band intensities to the standard ladder.

### Statistical Analysis

Data were analysed in Excel with one-way ANOVA with their respective p-values displayed on the figures. Error bars represent the sample standard deviation for triplicates.

## Results

### Alanine scanning of 11 amino acid tag yield enhanced variants

Given the short 11 amino acid, substituting amino acids could enhance the tag’s effectiveness or help identify key residues for further engineering to create an improved tag. To examine this, we designed a general expression cassette with NT11 as a tag at the N-terminal, followed by the protein of interest, and ending with an affinity tag for protein purification (see Figure 1A). The NT11 tag and its mutants would be combined with fusion partners, and the resulting protein yield after expression would indicate the overall effectiveness of mutagenesis compared to the wildtype.

**Figure 1.**
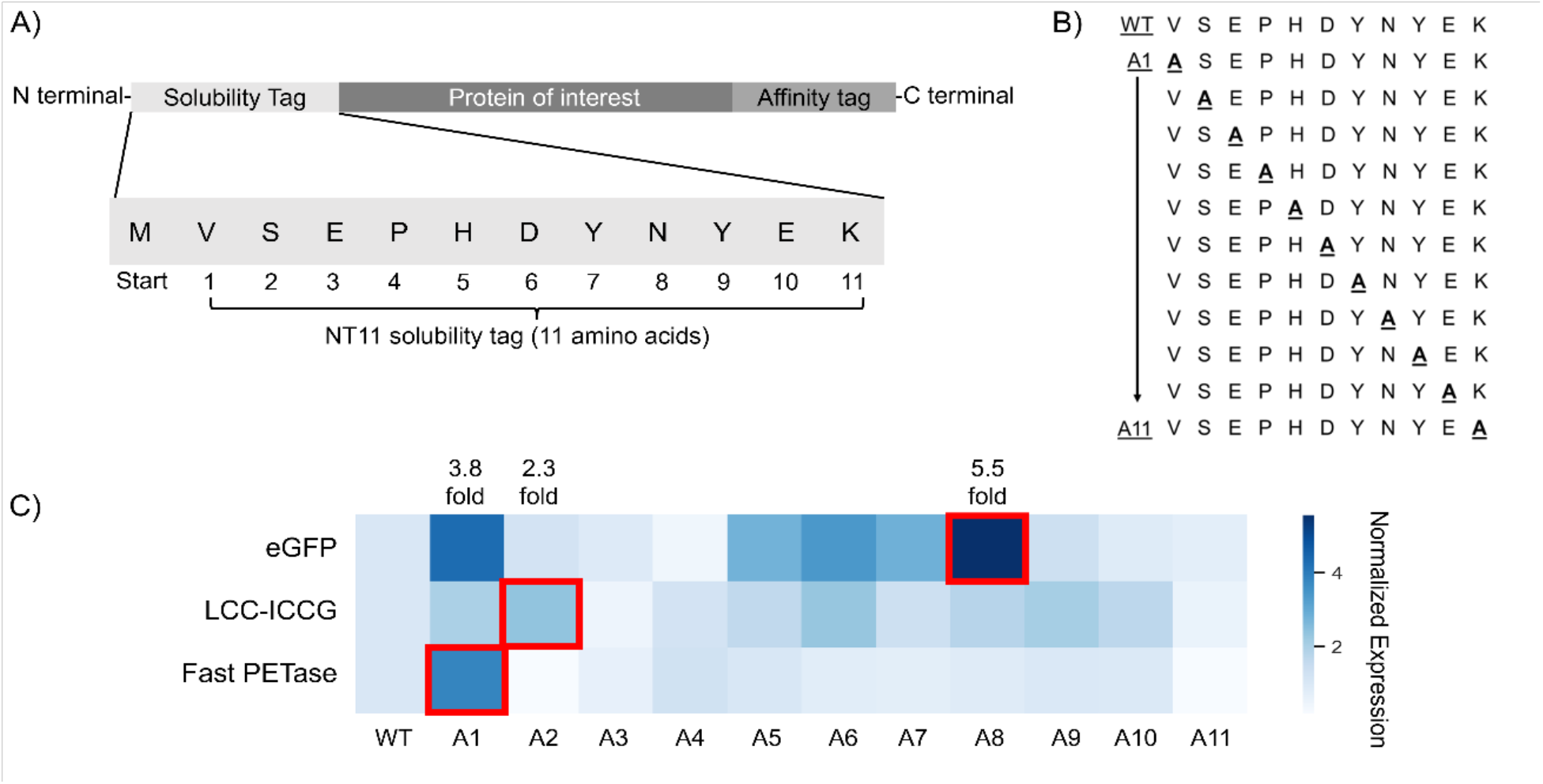
Alanine scan of the NT11 tag. A) General protein schema for protein expression B) Alanine scan: A total of 11 NT11 mutants were generated via single alanine substitutions along the entire length of the protein tag. C) Fold change in protein expression (densitometry) compared to wild type (WT) NT11 is represented as a heatmap featuring 3 proteins: enhanced green fluorescence protein (eGFP), leaf-branch compost cutinase, polyethylene terephthalate (PET) plastic degrading enzyme (Fast PETase). The highest expression for each protein is highlighted in red squares with its corresponding foldchange displayed above in the same column. The fold-change in graphical format can be found in supplementary Figure S1.

Single alanine substitution of each amino acid along the length of the NT11 tag produced 11 mutants, labelled NT11 A1 to A11 (Figure 1B). A total of 3 different proteins were paired with the NT11 tags and expressed to determine protein yield. The fluorescent protein eGFP (32.7kDa, pI 6.2, [18]) is a common reporter protein for determining protein production while engineered versions of a Leaf Compost Cutinase (LCC-ICCG, 31.5 kDa, pI 9.3, [19]) and PET degrading enzyme (Fast PETase, 27.6 kDa, pI 9.6, [20]) are both known plastic depolymerization enzymes important for their role in polyethylene terephthalate (PET) degradation and recycling. PET degrading enzymes are also known to have difficulty being expressed in *E. coli* with low reported yields, hypothesized to be due to the presence of two disulfide bridges [21-23].

From screening of single alanine mutated tags, it was observed that improved expression levels of up to 5 folds can be achieved with a single mutation, however these were dependent on the specific protein pairing (see Figure 1C). Even so, we observed that A1 position is consistently better with a 2 to 4-fold increase across the 3 proteins. The highest yields achieved were through different protein-tag pairings across eGFP, LCC-ICCG, Fast PETase (A8, A2 and A1 respectively-see Figure 1C). Besides improvement, certain mutants can also lead to decreased protein yields – including sites at positions A3 and A11 (see Figure 1D) where no or reduced soluble yield could be observed. Such residues correspond to the negatively charged glutamic acid and positively charged lysine respectively, suggesting the importance of having specific charges at their respective positions even in an 11-amino acid long peptide sequence.

In further work, the key residues can be further modified, going beyond alanine substitutions, to enhance the NT11 tag for specific protein pairings. However, these initial observations have demonstrated the effectiveness of a simple alanine scan as a rapid preliminary screen to improve yields for each unique protein, with 2 to 5-fold improvements compared to the wild-type tag.

### NT11 tag as a flexible enhancement motif

Upon performing *in silico* searches in Genbank using NT11 as a protein query, we discovered highly similar sequences (Figure 2A) present in a variety of natural proteins. These proteins include enzymes like mitochondrial transmembrane protein choline dehydrogenase and anti-pathogen acidic mammalian chitinase [24,25], as well as structural proteins like DNRLRE- domain containing proteins [26,27]. We found NT11-like sequences not only located near the N-terminal, as seen with dCA, but also close to the C-terminal in proteins such as acidic mammalian chitinase and DNRLRE-domain containing proteins. This discovery led us to hypothesize that the effect of the NT11 motif may not be confined to the prototypical N-terminus tag location.

**Figure 2.**
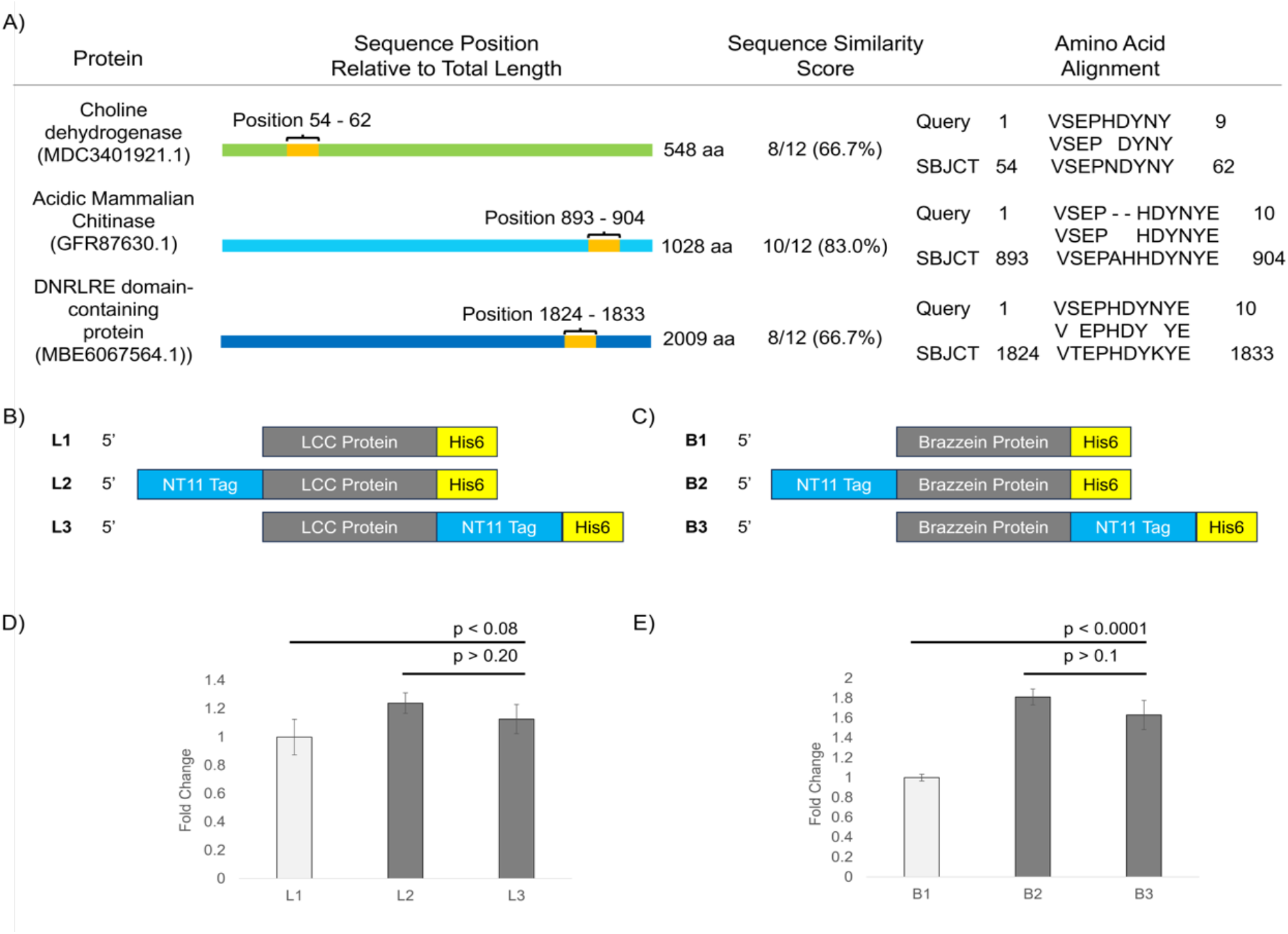
The flexibility of the NT11 tag and its ability to be placed on both ends of the protein while still functioning as an enhancement tag. A) Characterisation of amino acid sequences highly similar to NT11 were found across various naturally occurring proteins. B) and C) schema of the 3 protein constructs designed for each target protein (LCC-ICCG and brazzein). D) and E) Fold change of soluble protein yield with respect to the various NT11 placement, as calculated by densitometry. Experiments were performed with biological triplicates.

To explore the potential placement of NT11 at different termini of a recombinant protein, we designed three sets of constructs using two proteins: PET degrading enzyme LCC-ICCG (Figure 2B, [19]) and plant-derived sweet protein brazzein (Figure 2C, [28]). These proteins contain 2-4 disulfide bonds and are important proteins for applications of plastic recycling and alternative non-calorie sweeteners, where yields are important to achieving cost efficiencies.

The expression constructs of the proteins proceeded with (1) no tag, (2) NT11 placed at the N-terminal, and (3) NT11 tag placed at the C-terminal. Additionally, all constructs have a C-terminal His-tag to facilitate affinity purification. In both LCC-ICCG and brazzein, we observed similar trends in recombinant protein yield improvement with tags appended to either the N- or C-terminus of the target protein (see figure 2D and 2E). We have now demonstrated that the NT11 tag could be attached to either the N- or C-terminus of a paired protein, also bordered by affinity tags, while exploiting its yield-enhancing effects.

### Improved expression of growth factors

Here, using a combination of NT11 and NT11 A1 tags and location placement, we investigated the effects of incorporating NT11 to 2 structurally distinct growth factors, fibroblast growth factor 2 (FGF2) and human epidermal growth factor (hEGF). Thermostable FGF2-G3 [29] and hEGF [30] are 2 high value growth factors belonging to 2 separate growth factor families which garnered significant interest in recent years for their expression in prokaryotic systems in attempts to develop low costing, high yielding manufacturing routes [31,32]. Due to their high demand in cell culture work and as treatment components for injuries, there were constant need for new avenues of production [31,33]. A recently published work performed cost analysis of mitotic growth factors used in cell culture media formulations and found that currently, 1 mg of growth factor costs more than CAD$1000 (approx. USD$730 at time of writing) [32].

Although prokaryotes represent a potentially affordable source of growth factors, challenges such as protein aggregation, incorrect folding and general low yields has hindered their production [34]. The vast majority of current growth factor production in *E. coli* depend on refolding strategies after isolating the protein aggregates in the form of inclusion bodies, contributing to prohibitive costs [35]. Solubility tags were explored as a solution and worked to varying degrees, with some approaches requiring cumbersome purification processes or specialized equipment [35]. Some of the most successful designs recorded demonstrated the expression of functional purified FGF2 achieving 100 mg/L yields upon the removal of tag [36]. On the other hand, recent efforts at expressing high-yielding hEGF made use of continuous free-cell cultures aided by biofilm formation to achieve up to 52.4 mg/L yields [37], highlighting the need for simpler and more effective expression systems.

While the 2 growth factors differ in both size (∼20 kDa and ∼10 kDa for FGF2 and hEGF respectively) and their 3-dimensional structures (see Figure 3A, Figure S2 [38,39]), they are both small sized proteins (<20 kDa), making a small tag more suitable as a solubility enhancer. In this study, we investigated the effects of both location and mutation of NT11 on recombinant protein production. For each growth factor, we investigated the effects of incorporating NT11 tag in 5 constructs, namely (1) N-terminal tagged, (2) C-terminal tagged, (3,4) equivalent N and C-terminal tagged for NT11-A1 mutant and (5) NT11 A1 tagged at both the N and C-terminal (Figure 3B). Expression for these constructs was carried out in micro-bioreactors [40,41].

**Figure 3.**
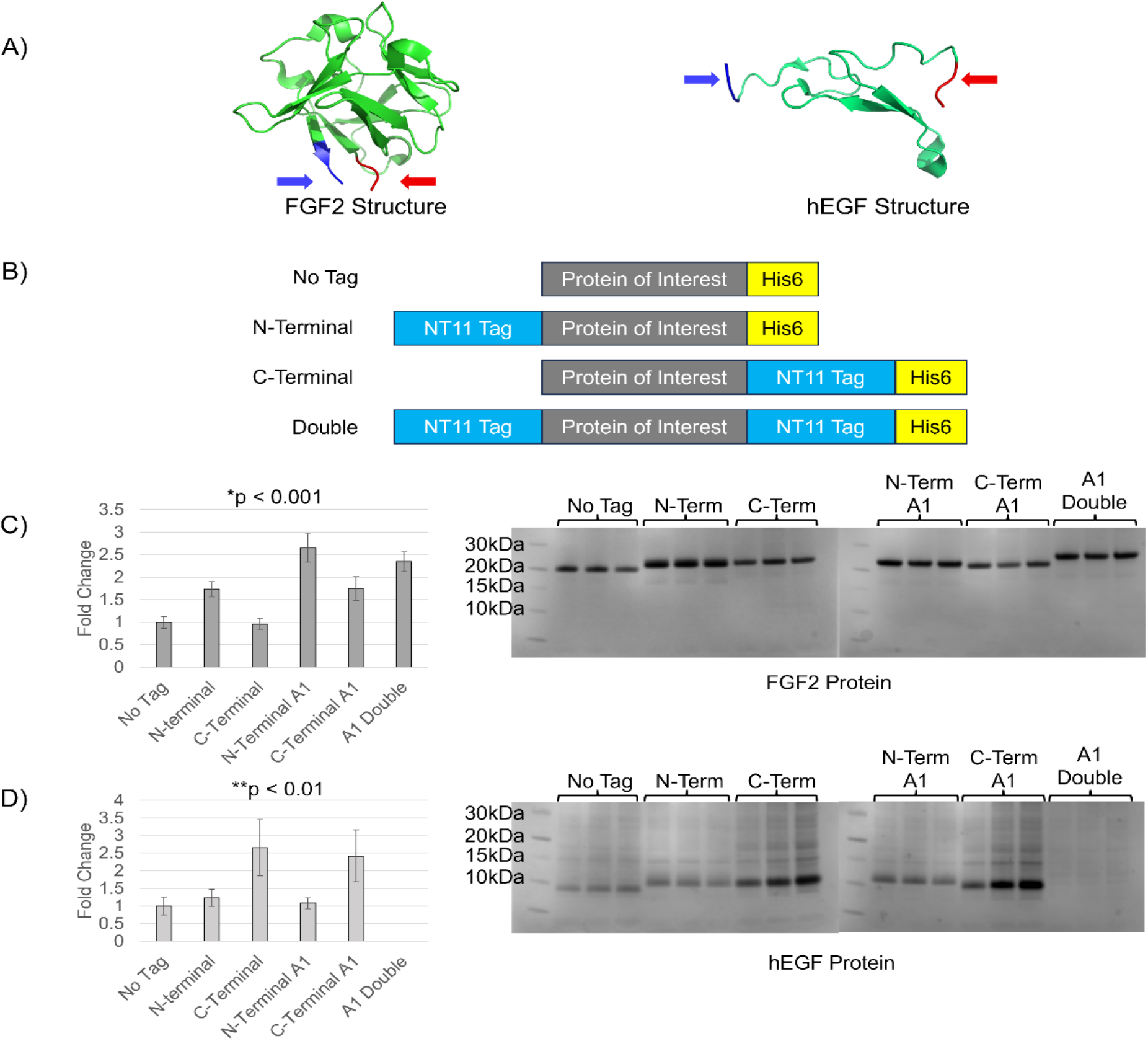
Growth factor expression under micro-bioreactor conditions. A) The 3-dimensional structure of FGF2 (PDB: 1BFB) [42] and hEGF (PDB: 1IXA) [43], with the N- and C- terminals indicated in blue and red respectively. B) The general schema of protein constructs, with both FGF2 and hEGF being targets for expression. C) and D) Comparison of fold change of tagged against non-tagged variants (left) and their respective SDS-PAGE gel of purified soluble proteins (right). Fold change of soluble protein yields were calculated based on densitometry. Experiments were performed in biological triplicates.

The purified growth factors showed distinct fold-change improvements over constructs with no NT11 tag. FGF2 protein demonstrated up to 2.6±0.3-fold yield improvement over the non-tagged construct while hEGF demonstrated up to 2.6±0.8-fold yield improvement (see Figure 3). Using the A1 mutant resulted in improved FGF2 performance, consistently outperforming non-mutant designs for both N- and C-terminal tagging. However, the A1 mutant tag had minimal impact on hEGF, suggesting the need for further optimization of the NT11 tag. The results demonstrate a preference for N-terminal tagging for FGF2 (Figure 3C) and C-terminal tagging for hEGF (Figure 3D). The different effects of tag mutagenesis and positional placement on these growth factors were likely contributed by their distinct structures. This also highlights that different tags are used depending on its circumstances. However, this new property of NT11 may allow for a more versatile tool for enhancement of recombinant protein production.

## Discussion

In this study, through further characterisation and engineering of NT11, we were able to expand upon the utility of a short peptide tag. To demonstrate its utility, we tested NT11 on a diverse range of proteins with varied surface charges, structures and disulfide bonds (Figure S2) [44], where we observed >60% improvements in all applications. Due to the short length of the NT11 tag (11 amino acids), the alanine scan was a quick method used to identify locations for potential optimization. It is important to note that these sites differed depending on the specific protein, as distinct fusion partners resulted in different sites. In future research, these sites can be further investigated to determine the most efficient amino acid substitutions.

In another optimization strategy, we have identified an additional property of NT11 based on our observations of similar NT11 sequences in nature. Here, we demonstrated the first examples of NT11 as an enhancer of recombinant protein yields at both the N- and C-termini in comparison to the protein of interest. This suggests that NT11, when combined with an affinity tag (such as a His-tag), can be positioned either before or after the protein of interest without impacting the enhancement capability of NT11. This flexibility opens up numerous possibilities for arranging protein-tags combinations, along with purification strategies.

With these new findings, we managed to double recombinant production of soluble growth factors in *E. coli*. FGF2 expression could be improved further using NT11, with mutant variant NT11 A1 being able to push expression above 2.5-fold. While novel methods have been employed for producing hEGF due to its low yielding nature in *E. coli* [37], we have shown that simply utilizing the NT11 tag was sufficient to double improve yield of recombinant protein expression. The preference for N- or C-terminal tagging for FGF2 and hEGF respectively would again emphasize on the context-specific importance of tag positioning while optimizing for fusion partners. Overall, the small size of NT11, coupled with its newly observed flexibility in placement and design, represent a valuable tag system for use in efficient recombinant protein expression and purification.

## Conclusions

Small solubility tags, such as NT11 tags, are highly attractive due its size and impact. In this study, we have further expanded on the functionality of this 11-amino acid tag. Through various optimisation methods such as alanine scan mutagenesis and N- or C-terminal positioning, the yields of recombinant protein fusion partners, ranging from enzymes to growth factors, could be improved >60%. Looking forward, the alanine scan is also a simple and effective initial optimization strategy for small tags, which can significantly improve recombinant protein yields.

## List of abbreviations

(*E. coli*): *Escherichia coli*
(FGF2): fibroblast growth factor 2
(hEGF): human epidermal growth factor
(SynIDPs): small synthetic intrinsically disordered proteins
(dCA): duplicated carbonic anhydrase
(LB): Luria Bertani
(IPTG): isopropyl-β-D-1-thiogalactopyranoside
(IMAC): immobilized metal affinity chromatography
(eGFP): enhanced green fluorescence protein
(LCC): leaf-branch compost cutinase.
(PET): Polyethylene terephthalate
(Fast PETase): plastic degrading enzyme

## Declarations

### Ethics approval and consent to participate

Not applicable

### Consent for publication

Not applicable

### Availability of data and materials

All data generated or analysed during this study are included in this published article [and its supplementary information files].

## Authors’ contributions

JB: conceptualization, methodology, investigation, formal analyses, writing-original draft, writing-review & editing. ET: Methodology, Investigation, writing-review & editing, ZWB: supervision, writing-review & editing, FTW: Supervision, conceptualization, methodology, investigation, formal analyses, writing-original draft, writing-review & editing.

All authors also read and approved the final manuscript.

## Acknowledgements

Not applicable

