## Supplementary material for "Further characterization and engineering of a 11-amino acid motif for enhancing recombinant protein expression": Figs S1-2, Tables S1-2

**Figure S1.** The protein expression fold change of each protein featured in the heatmap in Figure 1C presented in graphical format.

**Figure S2.** The overall surface charge of eGFP (PDB: 6YLQ), LCC-ICCG (PDB: 4EB0), FAST PETase (7SH6), Brazzein (4HE7), FGF2 (PDB: 1BFB), hEGF (PDB: 1IXA) at pH 7.0 and the visualization of protein surface charge with negative charge (red) and positive charge (blue).

**Table S1.** Table documenting the full amino acid sequence of the proteins expressed with wild type NT11 tag.

**Table S2.** Table documenting the primer sequences used for performing QuikChange™ PCR mutagenesis for creating the alanine scan library of plasmids.

### Supplementary data

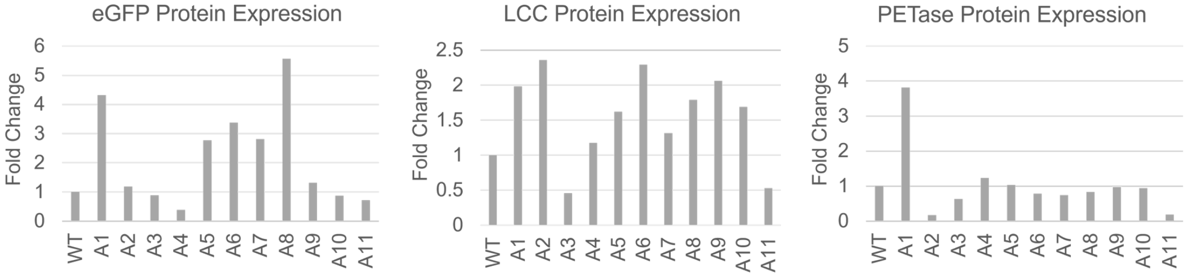


*Figure S2. The protein expression fold change of each protein featured in the heatmap in Figure 1C presented in graphical format. High-throughput single pass experiment with no repeats was performed.*

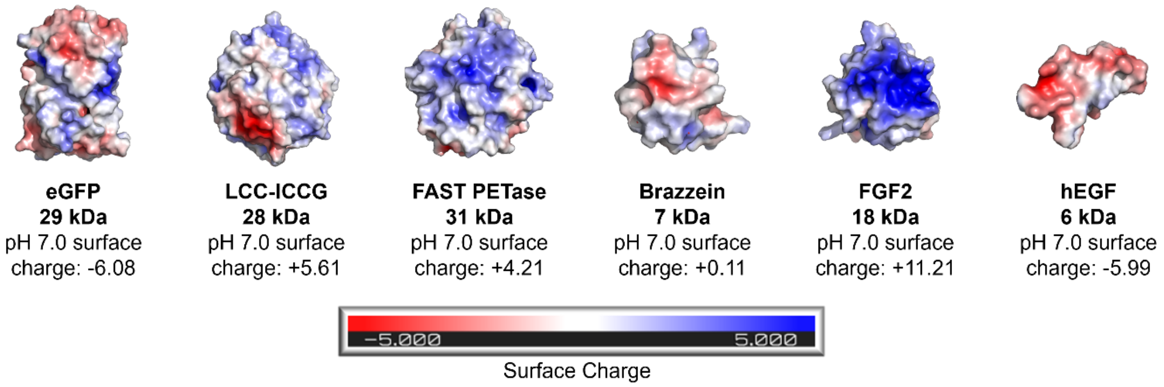


*Figure S3. The overall surface charge of eGFP (PDB: 6YLQ), LCC-ICCG (PDB: 4EB0), FAST PETase (7SH6), Brazzein (4HE7), FGF2 (PDB: 1BFB), hEGF (PDB: 1IXA) at pH 7.0 and the visualization of protein surface charge with negative charge (red) and positive charge (blue).*

*Table S1. Table documenting the full amino acid sequence of the proteins expressed with wild type NT11 tag.*

| | Amino Acid Sequence | |
|---|---|---|
| Plasmid ID | Tag Sequence | Target protein sequence |
| pET28-NT11-eGFP | VSEPHDYNYEK | VSKGEELFTGVVPILVELDGDVNGHKFSVSGEGEGDATYGK<br>LTLKFICTTGKLPVPWPTLVTTLTYGVQCFSRYPDHMKQHD<br>FFKSAMPEGYVQERTIFFKDDGNYKTRAEVKFEGDTLVNRI<br>ELKGIDFKEDGNILGHKLEYNYNSHNVYIMADKQKNGIKVNF<br>KIRHNIEDGSVQLADHYQQNTPIGDGPVLLPDNHYLSTQSAL<br>SKDPNEKRDHMVLLEFVTAAGITLGMDELYK |
| pET28-NT11-LCC | VSEPHDYNYEK | MSNPYQRGPNPTRSALTADGPFSVATYTVSRLSVSGFGGG<br>VIYYPTGTSLTFGGIAMSPGYTADASSLAWLGRRLASHGFV<br>VLVINTNSRFDGPDSRASQLSAALNYLRTSSPSAVRARLDA<br>NRLAVAGHSMGGGGTLRIAEQNPSLKAAVPLTPWHTDKTF<br>NTSVPVLIVGAEADTVAPVSQHAIPFYQNLPSTTPKVYVELC<br>NASHIAPNSNNAAISVYTISWMKLWVDNDTRYRQFLCNVND<br>PALCDFRTNNRHCQ |
| pET28-NT11-FAST_PETase | VSEPHDYNYEK | MNFPRASRLMQAAVLGGLMAVSAAATAQTNPYARGPNPTA<br>ASLEASAGPFTVRSFTVSRPSGYGAGTVYYPTNAGGTVGAI<br>AIVPGYTARQSSIKWWGPRLASHGFVVITIDTNSTLDQPESR<br>SSQQMAALRQVASLNGTSSSPIYGKVDTARMGVMGHSMG<br>GGGSLISAANNPSLKAAAPQAPWHSSTNFSSVTVPTLIFACE<br>NDSIAPVNSSALPIYDSMSQNAKQFLEIKGGSHFCANSGNS<br>NQALIGKKGVAWMKRFMDNDTRYSTFACENPNSTAVSDFR<br>TANCS |
| pET28-NT11-Brazzein | VSEPHDYNYEK | MQDKCKKVYENYPVSKCQLANQCNYDCKLDKHARSGECF<br>YDEKRNLQCICDYCEYP |
| pET28-NT11-FGF2 | VSEPHDYNYEK | PALPEDGGSGAFPPGHFKDPKLLYCKNGGFFLRIHPDGRVD<br>GTRDKSDPFIKLQLQAEERGVVSIKGVCANRYLAMKEDGRL<br>YAIKNVTDECFFFERLEENNYNTYRSRKYPSWYVALKRTGQ<br>YKLGSKTGPGQKAILFLPMSAKS |
| pET28-NT11-hEGF | VSEPHDYNYEK | NSDSECPLSHDGYCLHDGVCMYIEALDKYACNCVVGYIGER<br>CQYRDLKWWELR |

*Table S2. Table documenting the primer sequences used for performing QuikChange™ PCR mutagenesis for creating the alanine scan library of plasmids.*

| Plasmid ID | Primer Nucleotide Sequence (5' to 3') |
|---|---|
| pET28-p538-NT11-A1 | Forward: TGCGGTTCGGACGCcatGGTATATCTCCTT<br>Reverse: AAGGAGATATACCatgGCGTCCGAACCGCA |
| pET28-p538-NT11-A2 | Forward: ATAATCATGCGGTTCCGCTACcatGGTATATCTCC<br>Reverse: GGAGATATACCatgGTAGCGGAACCGCATGATTAT |
| pET28-p538-NT11-A3 | Forward: AGTTATAATCATGCGGCGCGGATACcatGGT<br>Reverse: ACCatgGTATCCGCGCCGCATGATTATAACT |
| pET28-p538-NT11-A4 | Forward: CATAGTTATAATCATGCGCTTCGGATACcatGG<br>Reverse: CCatgGTATCCGAAGCGCATGATTATAACTATG |
| pET28-p538-NT11-A5 | Forward: TGCCTTCTCATAGTTATAATCcgcCGGTTCGGATACca<br>Reverse: tgGTATCCGAACCGGCGGATTATAACTATGAGAAGGCA |
| pET28-p538-NT11-A6 | Forward: TGCCTTCTCATAGTTATAcgcATGCGGTTCG<br>Reverse: CGAACCGCATgcgTATAACTATGAGAAGGCA |
| pET28-p538-NT11-A7 | Forward: CTGCCTTCTCATAGTTcgcATCATGCGGTTCG<br>Reverse: CGAACCGCATGATgcgAACTATGAGAAGGCAG |
| pET28-p538-NT11-A8 | Forward: CCTGCCTTCTCATAcgcATAATCATGCGGTTC<br>Reverse: GAACCGCATGATTATgcgTATGAGAAGGCAGG |
| pET28-p538-NT11-A9 | Forward: CTCCTGCCTTCTCcgcGTTATAATCATGCGGT<br>Reverse: ACCGCATGATTATAACgcgGAGAAGGCAGGAG |
| pET28-p538-NT11-A10 | Forward: GCTCCTGCCTTcgcATAGTTATAATCATGCG<br>Reverse: CGCATGATTATAACTATgcgAAGGCAGGAGC |
| pET28-p538-NT11-A11 | Forward: cAGCTCCTGCcgcCTCATAGTTATAATCATGC<br>Reverse: GCATGATTATAACTATGAGgcgGCAGGAGCTg |
